# Single Cells of *Pseudomonas aeruginosa* Exhibit Electrotaxis and Durotaxis Behaviours

**DOI:** 10.1101/020511

**Authors:** Lee Preiss, Suresh Neethirajan

## Abstract

*Pseudomonas aeruginosa* is frequently associated with nosocomial infections, including polymicrobial wound infections and the complex biofilm communities that reside within the cystic fibrosis lung. *P. aeruginosa* utilizes flagellum-mediated motility to approach, attach to, and spread across a surface using a combination of swimming, swarming and twitching (type IV pili extension and retraction) motility. We report an innovative approach to study the motility of single *P. aeruginosa* cells in microfluidic channels possessing different structural geometry, all with the flexibility of being able to manipulate stiffness gradients and electric fields to investigate changes in motility in response to specific stimuli. *P. aeruginosa* cells exhibit restricted motility in reduced microchannel spaces, with preferential migration toward a stiffer region in a rigidness gradient of substrate medium and preferential migration toward a positive electrode in presence of a pulsed or successive electric field. This single-cue environmental study using microfluidic technology lays the groundwork for multi-cue experimentation to more faithfully mimic conditions *in vivo*, demonstrating just some of the advantages of this technique. This study is designed to examine the interplay between surface rigidity, mechanical, and electrical cues to pave the way for improvements in the design of anti-fouling surfaces for biomedical applications and to identify new ways to inhibit bacterial biofilm growth through motility restriction.

## Introduction

Wound matrices are complex structures with heterogeneous pore spaces, such that the concentration of chemicals, nutrients, and enzymes essential for bacterial survival may vary significantly across the micrometric spatial scale (Lawrence and Madihally, 2008). The porosity and the tortuosity characteristics inside the wound matrix therefore influence bacterial motility (Wiegand and White, 2013). Topography and surface chemistry also influence bacterial motility and may ultimately be a deciding factor in the decision to retain motility or attach (Belas, 2014; Romling, 2013; Costerton et al., 1999). The inherent spatial heterogeneity of wound matrices and medical implant tissue surfaces, along with the dynamic fluctuations that occur in the chemical gradients and hydration conditions of the wound microenvironment, significantly affects the responses of pathogenic bacteria. Of these wound characteristics, surface roughness greatly impacts bacterial attachment and additional hydrodynamic factors that collectively help to shape bacterial motility (Perera-Costa et al., 2014). To better understand the impact of diffusional heterogeneity within the wound matrix, we explored the influence of surface features on the motility of individual *Pseudomonas aeruginosa* cells.

Understanding the motility of single *P. aeruginosa* cells will likely provide insight into the design of new antifouling surfaces for food processing and biomedical applications, as *P. aeruginosa* is a well characterized biofilm-forming pathogen that survives on abiotic surfaces and within the human host (Hall-Stoodley and Stoodley, 2009). Understanding changes in *P. aeruginosa* motility in response to specific surface features, such as stiffness, will provide a better understanding necessary to develop strategies to impede bacterial movement during the treatment/prevention of bacterial biofilm infections. In support of this approach, it has been shown that the local topography of cell culture surfaces affects cell directional persistence and motility (in mammalian cells). For example, localized topographical features on PDMS substrates can induce an abrupt loss in the directional persistence of Osteoblast MC3T3-ET cells (Tang et al., 2014).

Mechanotaxis and durotaxis behaviours are strongly influenced by the external environment (Mott, 2014; Mitchell and Kogure, 2006). For example, microfluidic analysis of bovine aortic cells shows that the aortic cells preferentially migrate from substrates of lesser stiffness to surfaces of higher stiffness (Isenberg et al., 2009). The response to specific stimuli can also induce physiological changes in *bacterial cells*, although this is less well understood than in eukaryotic cells. It has been established that pathogenic bacterial species use several mechanisms to move and attach to surfaces, including electrochemical attraction and directed motility (Dunne, 2002). Through a better grasp of the motility exhibited by bacterial cells in response to environmental cues, we may be able to develop strategies to favorably alter motility and ultimately discourage biofilm formation.

Electrotherapy approaches have steadily gained interest as potential novel strategies to prevent bacterial biofilm infection and ultimately promote wound healing (Birkenhauer and Neethirajan, 2014; Zhao et al., 2006). Electric stimulation plays a significant role in regeneration, morphogenesis, and wound healing (McCaig et al., 2009), and electric fields can be used to control and direct cellular migration via changes in shear flow and chemotaxis (Gov, 2014). A significant improvement in wound healing might be obtained through the implementation of strategies that employ/develop novel bioelectric wound dressings. To gain critical information for the development of such technologies, we need to first investigate the response of individual pathogenic bacterial cells to electric fields and fluctuations in these fields. Electrotactic cues do not exert a bulk force on epithelial cells and act downstream of motility pathways (Cohen et al., 2014). Current density drives the cellular motility, not the electric field itself (Cohen et al., 2014). Electrotaxis has been studied extensively in epithelial cells and keratinocytes, but has not been explored in individual bacterial cells. Similarly, osmolarity and bacterial chemotaxis have been investigated using microfluidic platforms, while durotaxis has not been studied thoroughly.

Microfluidic platforms are ideal for the study of single cell electrotaxis due to their flexible design, enhanced modeling accuracy, and their ability to control and manipulate the cellular environment (Neethirajan, et al., 2011; Rusconi et al., 2014). Microfluidic technology is a tool that may enhance our systematic understanding of complex biophysical and biochemical processes involved in bacterial cell motility on the scale of single cell interactions (Hol and Dekker, 2014). Microfluidic approaches offer several advantages, including reduced sample volume, reduced reagent cost, and scalability to high throughput screening applications, ultimately streamlining complex assay protocols and better emulating the biological microenvironment (Sackmann et al., 2014).

Although eukaryotic cells have been shown to respond to a variety of mechanical cues (Discher et al., 2005; Kuo et al., 2012), mechanotaxis and electrotaxis have not been extensively studied in pathogenic bacterial species. The mechanisms of mechanical force sensing in bacterial are largely unknown (Mendez-Vilas, 2010). Using microfluidics, we begin to investigate the complex mechanisms that govern microbial cell migration in a biofilm environment. The overall goal of this study was to understand: (1) the effect of surface structures on the swimming dynamics and motility of *Pseudomonas aeruginosa* by incorporating relevant wound materials (e.g. collagen), and (2) the electrotaxis and durotaxis behaviours of single *P. aeruginosa* cells using microfluidic platforms.

## Experimental Procedure

### Microfluidic Devices Design and Preparation

A silicon and SU-8 master template of the microfluidic device was used to fabricate single-use PDMS devices for experimentation. The master template was first subjected to a silane bath under negative pressure in a desiccator for a minimum of 6 hours. A 30:3 wt/wt (10:1) mixture of PDMS and its associated binding agent was mixed thoroughly and dispersed evenly on the device. The device with the liquid PDMS was degassed and heated to 60°C for 4 hours to ensure complete polymerization. The mold was then removed from the silicon master template and cut such that its dimensions did not exceed those of the glass slide. Holes were punched at the inlets and outlets of the device from the bottom side. The device and slide were cleaned with tape to remove debris, followed by plasma cleaning for 1 minute to functionalize the bonding surfaces. Finally, the device was adhered to the slide using minimal pressure and reheated to 60°C for 30 minutes. In total, 5 microfluidic devices were used throughout the experimental study. Three devices were used for microchannel geometry trials (Figures 1A, 1B, and 1C), a single serpentine channel (Figure 1D) was used for the durotactic observation, and a Y-Channel microfluidic device (not shown) was used for the electrotactic study. The gradient generating device (Figure 1D) utilized for the durotaxis experiments contains 8 rows of mixing modules allowing for 10 different concentration ratios of the inlet solutions to flow into the observation module (DiCicco, 2013). The device design includes two sections, consisting of a mixing module and an observation module with the following dimension details: 1) Mixing Module: Eight rows of mixers with micro-channels of 20 micron in depth, 100 microns in width and a total length of 18,750 microns in length (across all 8 rows). 2) Observation Module: Where the individual bacterial motility is observed, it is 1000 µmin width, 40 µm in depth, and 12,000 µm in length (between the inlet and outlet). The design configuration of the Y-channel device (not shown) for electrotaxis experiments had 3 inlets and 1 outlet with 800 μm wide, 125 μm deep central observation channel with a reservoir volume of 100 μL.

**FIGURE 1.**
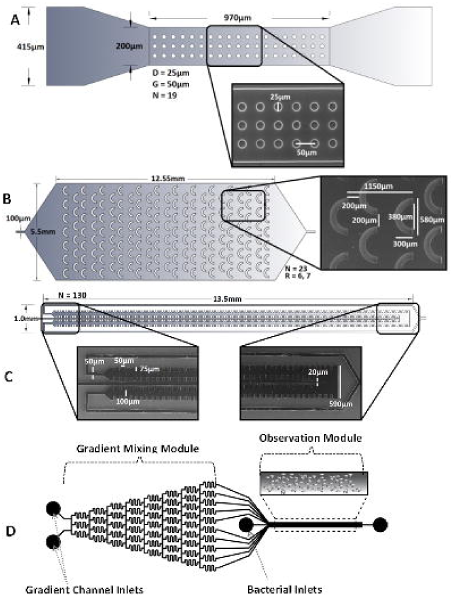
**S**canning electron microscope (SEM) images of the **A)** single-channel circular deformation, **B)** semi-circular shell, and **C)** square devices used in the micro-channel geometry study, **D)** Gradient-mixing module (20 x 100 x 18750 μm) and a chemotaxis observation module (20 (depth) x 1000 (width) x 11500 (length) μm). The widths of the two gradient channel inlets entering the device are 500 μm and the width of the bacteria inlet is 50 μm.

Microfluidic platforms designs were created using AutoCAD software and fabricated by following the standard soft lithography protocol as detailed by Qin et al. (2010). The layout and the device design dimensions and configurations of the microfluidic platforms are shown in Figures 1A, 1B, 1C and 1D.

Upon fabrication of the template, the devices were ultimately etched onto a thick photoresist wafers by using e-beam photolithography. Upon prebaking, photomasks with the designed microchannel geometries were placed onto the coated silicon wafers and exposed to UV using a UV exposure system (UV-KUB, Kloé, France). After the post-baking and developing, each microfluidic platform was used as a mold for creating the PDMS chip. A 10:1 (w/w) mixture of PDMS polymer and the curing agent (Sylgard, Dow Corning, Burlington, ON, Canada) was stirred thoroughly and degassed under vacuum. Then the mixture was poured onto the master mold and cured at 75°C for 4 h. After curing, the PDMS replica was peeled off from the master, punched with holes to provide inlets and outlet and bonded onto a glass slide (25 × 75 × 1 mm, VWR International, Suwanee, GA, USA) after oxygen plasma treatment for 40 s.

### Cell Culture

The strain tested in the present study was a clinical isolate of *Pseudomonas aeruginosa (Bk-76),* which was derived from the canine ear skin infections. The bacteria were seeded on Tryptic Soy Agar (TSA) plates and incubated for 24 hours at 37°C, followed by storage at 4°C until further use. Fresh plates were prepared every two weeks. Individual colonies were cultured in 5 ml Bacto Tryptic Soy Broth (TSB) for 16 hours with continuous shaking at 140 rpm and 37°C. A 1 ml sample of the cell suspension was subjected to centrifugation at 3000 rpm for 3 minutes. The supernatant was discarded and this step was repeated with additional 1 ml of distilled water. The cell pellet was resuspended in 1 ml of chemotactic buffer (CB). The chemotactic motility buffer was prepared by mixing 11.2 g of potassium phosphate dibasic, 4.8 g of potassium phosphate monobasic, 2 g of ammonium sulphate, 0.25 g of magnesium sulphate and 1 g of polyvinylpyrrolidone (Sigma Aldrich, St. Louis, MO) in 1 litre of deionized water.

### Microscopy

The microfluidic platform was placed on the micrometre stage of the Nikon Eclipse Ti inverted microscope, and the single cell resolution images and videos were captured using a Nikon Ds-Qi microscope camera and Nikon NIS Elements BR version 4.13 software (Nikon Instrument Inc., Melville, NY). Microscope hardware settings consisted of an S Plan Fluor 40× objective in the phase contrast 1 mode, with the NCB and D filters on. The video was captured at 640 × 480 resolution in a 40 ms exposure, with 2× analog gain and at 15 fps in 1 minute-long sections.

### Microchannel Geometries

Three different microfluidic devices with varying geometries (circular, semi-circular shell, and square cut outs; Fig. 1A, 1B and 1C, respectively) were selected for this study. All devices were prepared in the same fashion. The devices were initially sterilized with 70% isopropyl alcohol and rinsed with distilled water. They were then coated with 0.1% sterile filtered bovine serum albumin (BSA) at room temperature for 10 minutes, with sealed inlets and outlets. After incubation, the devices were rinsed with distilled water. Cultured bacterial cells were then injected into this environment, and the inlet and outlet were resealed to allow for fluid settling over 15 minutes.

### Durotaxis

The serpentine gradient generating microfluidic device (Fig. 1D) was used to prepare a linear stiffness gradient of Type-1 Collagen. Collagen at the desired concentration was placed in one of the inlets and a control salt solution (PBS) was placed in the other. The solutions were initially injected at 100 μL/h flow rate to fill the serpentine channels and remove any air bubbles in the device and then reduced to 20 μL/h flow rate for proper mixing of the observation module for 15 minutes. The device inlets were then sealed, and the chambers were placed in an incubator for 30 minutes at 37°C for the formation of a collagen gel. Thereafter, the inlets were injected with PBS to wash excess collagen from the channels. The PBS solution was maintained throughout the experiment at 5 μL/h flow rate to prevent the backflow of bacteria. The cell pellet was stained with Syto-9 Live/Dead dye (Life Technologies Inc, Burlington, Canada) before resuspension in chemotactic buffer. To make the dye solution, 3 μL of dimethyl sulfoxide with 1.67 mM SYTO 9 dye and 1.67 mM propidium iodide was suspended in 1 ml water. Two hundred μL of this dye solution was then added to the washed bacterial cells (suspended in 1 ml of water) to yield 1.2 ml of dyed cell solution. The staining solution was combined with the cell pellet and mixed thoroughly and kept in the dark for 30 minutes. 1 ml of chemotactic buffer was then added followed by the injection of the cell suspension into the cell inlet of the microfluidic device at a flow rate of 20 μL/h. The microscope and viewing stage were protected from exposure to light. Bacterial cells were viewed with the EGFP microscope filter with a halogen light. Time lapse observations were made every minute for 30 minutes at a 640 × 480 resolution with 2 × 2 binning and 1.28 × analog gain. The view was centered about the middle of the observation module at a magnification where the width of the whole channel was visible.

### Electrotaxis

A Y-Channel microfluidic device was used for all electrotactic experiments where the outer inlets were plugged for all trials. The device was sterilized with 70% IPA and rinsed with distilled water. Bacterial cells were then injected into the device suspended in conductive chemotactic buffer medium. A platinum electrode was then inserted into both the inlet and the outlet of the device and sealed with paraffin wax. The fluid was allowed to settle over the course of 15 minutes before any data was recorded. The positive and the negative terminals of a variable DC power supply were connected to each electrode and we began to record observations. A 30-second video was recorded for cell motility at 0, 5, and 10 minutes after application of each voltage setting. Data was acquired at intervals of 0.0 V, 0.5 V, 1.0 V, 1.5 V, and 2.0 V applied in a successive manner. Data was then also recorded in fresh trials with 1.0 and 2.0 V applied without exposure to previous voltages.

### Cell Motility and Data Analysis

The video data recorded from the microchannel geometry trials and electrotaxis trials were processed in a similar manner. The videos were cut into 15 second fragments and analyzed with ImageJ software. The videos were adjusted to subtract the background, averaging the grayscale intensity of the full stack and removing it from the video. The video was then processed into binary format to clearly view cell motility. The plugin Manual Tracker was used to record cell motility for 50 bacterial cells per device in the micromechanical trials, and 30 bacterial cells per voltage and time step in the electrotaxis trials. The data files were then imported into the Ibidi Chemotaxis and Migration V2.0 software (Munich, Germany) for motility analysis.

The durotaxis time-lapse data was also analyzed with ImageJ software. The stacks were imported into the program and analyzed by creating a profile plot of fluorescent intensity. The images were thresholded with the Otsu-thresholding technique and fully selected for an intensity measurement. Data were collected from the start of the time lapse (t = [0, 5min]) until the end (t = [25, 30min]) to view a trend in bacterial movement. Phase-contrast images of cells on various regions of the collagen stiffness pattern across the channel were acquired using a 10X objective lens. Images were analyzed using ImageJ software (NIH) to subtract the background and to measure the cell area and the trajectories of the centroids of the cells along the stiffness gradient. Additional details regarding the methodology for calculating the different migration indexes can be found elsewhere at Zengel et al., (2011). Rayleigh test was used to test whether the deviations of the cell directions from the direction of electric field were significant.

### Stiffness Measurement Using AFM

Atomic Force Microscopy was performed using an Agilent 5500 series ILM-AFM under ambient conditions. The stiffness gradient across the channel was verified using the indentation measurement conducted in force mode. An area of 20 × 20 µm in the middle of each collagen pattenr with a 4 × 4 indentation array with 20 µm spacing was probed from the glass slide and analysed using the protocol of Costa (2006).

### Statistical Analysis

Quantitative data for cell motility behaviour were presented as the mean ± standard error. One-way ANOVA tests were also employed to determine the statistical significance of the differences between the two or more groups of the studied samples. Statistical analyses were performed using Origin Pro software (Origin Lab, Northampton, MA). A significance value of α = 0.05 was used for all tests. For cell migration across stiffness gradient, experiments were repeated three times (n=3) to acquire time lapsed images, and were segmented for cell positions using ImageJ software.

## Results

### Surface geometry alters chemotaxis parameters

We analyzed several single cell motility characteristics in their individual single-cue environments. The key traits that we characterized were cellular velocity, forward migration coefficients (FMI), the center of mass (CoM), and directness. The xFMI and yFMI represent the efficiency of the forward migration of cells, in relation to the x- or y- axis. The larger the index on an axis, the stronger the chemotactic effect is on this axis. For simplification, it is assumed that either the x-axis or the y-axis is parallel to the direction of the chemotactic gradient. The center of mass (CoM) represents the averaged point of all cell endpoints. Its × and y values indicate the direction in which the group of cells primarily traveled. The directionality is calculated by comparing Euclidean and accumulated distance. It is a measure of the directness of the cell trajectories. A directionality of D → 1 equals a straight-line migration from start to endpoint.

The microchannel geometries varied from a restricted circular deformation (Fig. 1A) to a semi-circle shell (Fig. 1B) to a square-tooth object (Fig. 1C). If the center of mass was mapped to the origin, we concluded that there was no flow (Table 1). The velocities of individual *P. aeruginosa* cells did show some minor fluctuations, based on their mechanical geometries and spatial restrictions. The circular microfluidic device (Fig. 1B) was restricted to a shallow depth of 10 μm with smaller boundaries. We observed a significant drop in the velocity in this device to 11.02 μm/s, when compared to 19.87 μm/s and 20.18 μm/s for semi-circular and square-tooth geometries, respectively (Fig. 2). Although cell directness was not statistically different among the different chamber geometries, the value was the highest in the smallest spatially-restricted microfluidic device (circular geometry; 0.138) compared to 0.100 and 0.091 for the semi-circular shell and sqre-tooth devices, respectively (Table 1). Other than reducing the run-trajectories of an individual bacterium, the values did not follow any specific trend when studied under different geometries with immediate effects on directness.

**TABLE 1.** Motility characteristics of single cell. ***Pseudomonas aeruginosa* on micropatterned surfaces inside microfluidic platforms of various designs. Forward migration coefficients and center of mass show a reputable environment for the study. The directness is comparable in all three microfluidic devices. Values are mean based on a population of 50 cells for 60 second tracking.**

| Microfluidic Device |  |  |  |
| --- | --- | --- | --- |
|  | Circular | Semi-circular shell | Square-Tooth |
| x FMI | 0.018 | -0.008 | -0.0111 |
| y FMI | 0.008 | 0.002 | 0.0030 |
| x CoM ($\mu\text{M}$) | 1.631 | -0.833 | -2.609 |
| y CoM ($\mu\text{M}$) | 0.980 | -1.768 | 0.829 |
| Directness | 0.138 | 0.1002 | 0.0909 |

**FIGURE 2.**
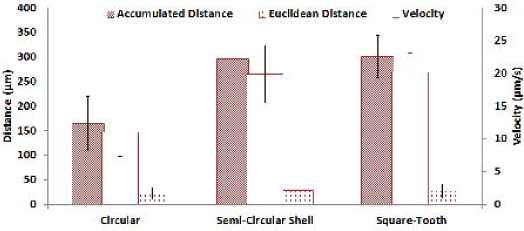
Micro-channel geometry effect on single-cell accumulated distance, Euclidean distance and velocity when subjected to differing geometries (circular, semi-circular shell and square-tooth). Euclidean distances were comparable but a significant drop was noted in velocity and accumulated distance in the circular microfluidic device due to spatial restriction.

### *P. aeruginosa* is attracted to environments of increasing stiffness

The assessment of *P. aeruginosa’s* response to greater rigidness, modeled by increasing the concentration of wound-relevant type-1collagen, is depicted in Fig. 4. With a slow flow rate of 20 μl/hr, bacterial movement exhibited little dependency on time (Fig. 4). Fluorescent bacteria shifted towards the left axis (increasing stiffness) of the observation module when observed over a 30 min block of time. Collagen is the most abundant protein in the human body and the major component of connective tissue. Aminoacids namely glycine, proline and hydroxyproline are present in collagen. The polymerization of collagen is a thermal process at approximately pH>6.5 which is compatible with the pH of the animal tissue (Nair and Laurencin, 2007). The scaffolds made of collagen inside the tissue forms adhesion sites for the bacterial cells. The results of AFM indentation analysis confirmed that there is stiffness gradient across the width of the observation channel of the microfluidic gradient generator and it ranged from 2 kPa to 70 kPa over a distance of 1000 µm (Fig. 3). Additional surface roughness measurement using contact mode AFM imaging revealed that the surface had a roughness of 70 nm. The recorded bacterial cell migration over 20 h showed that cells were initially interacting with the stiffness boundary, and the bacterial cell trajectories of the centroids were along the gradient direction. Image analysis results showed that the Pseudomonas aeruginosa cell moved from the softer regions to the stiffer regions clearly indicating that the bacterial cells were preferentially migrating towards stiffer regions.

**FIGURE 3.**
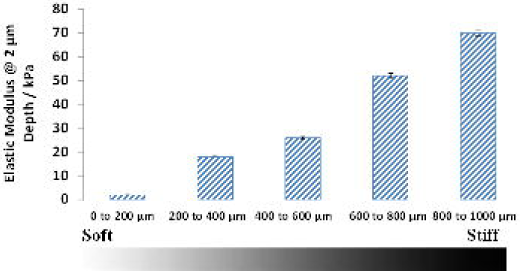
Verification of stiffness gradient inside the microfluidic serpentine channel platform. Apparent elastic moduli of the collagen gradient measured by AFM indentation (n=5).

**FIGURE 4.**
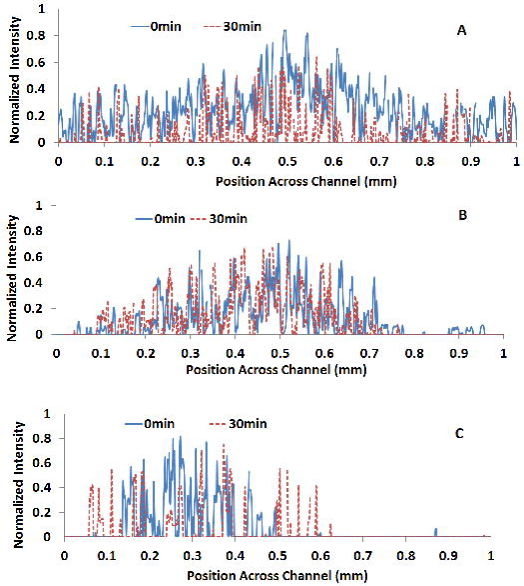
*Pseudomonas aeruginosa* cell migration on stiffness gradient. Intensity profiles over channel position of fluorescent bacteria with Control (A), 0.75 μg/ml (B) and 150 μg/ml (C) type-1 collagen used for stiffness gradient generation.

### Application of current impacts bacterial motility

The application of DC voltage to a microfluidic-controlled environment through chemotactic buffer resulted in altered cell motility (Fig. 5). The microfluidic channel filled with the buffer solution in all electrotaxis experiments showed a resistance of ∼0.84 MΩ. The accumulated distance traveled by individual bacterial cells was inversely proportional to the duration of the applied voltage. Single-cell velocities were also reduced from 18.82 μm/s (0.0V, t = 0) to 14.73 μm/s (2.0V, t = 10). Small changes occurred with each successive voltage application and became more obvious as the time of voltage application was extended (Fig. 5). This trend is apparent at the 1.5 V applications, as cell velocity drops from 17.75μm/s to 16.60μm/s to 15.01μm/s at time intervals of 0, 5, and 10 minutes, respectively. The Euclidean distances for individual bacterial cells did not show any changes, nor did they follow any specific trend over the course of successive voltage applications. Any differences observed were determined to not be statistically significant (P > 0.05). However, the cell directness was greatly affected (P ≤ 0.05) (Fig. 5) with voltage application, accompanied by changes in the center of mass. Directness increased from 0.145 to 0.201 when comparing the voltages of 0.0 V at t = 0 to 2.0 V at t = 10. The directness was dramatically increased when the bacterial cells were subjected to prolonged exposure to an applied electric field. The center of mass was also of importance in this situation, as it became increasingly positive, as the voltages were sustained for longer time periods. The yCOM values, which represent the overall movement of bacterial cells toward the positive electrode, increased to 26.48 μm when a 2.0 V current was applied for 10 min. This is a significant change compared to the original values that ranged from -8.15 to 0.204 before the application of an electric field (Table 2). Therefore, with an applied electric field, it can be expected that individual bacterial cells will move towards the positive electrode, representing a more direct trajectory path, as measured by directness.

**FIGURE 5.**
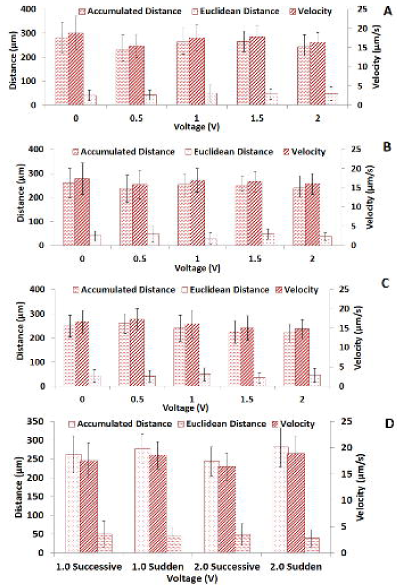
Successive voltage comparison on *P. aeruginosa* when subjected to 0.0, 0.5, 1.0, 1.5 and 2.0V at three time intervals of **A)** 0, **B)** 5, and **C)** 10 minutes. (D) Successive vs. sudden voltages applied of the same magnitude to note any major difference in application method. Both 1.0 and 2.0 V were tested at t = 0 min.

**TABLE 2.**
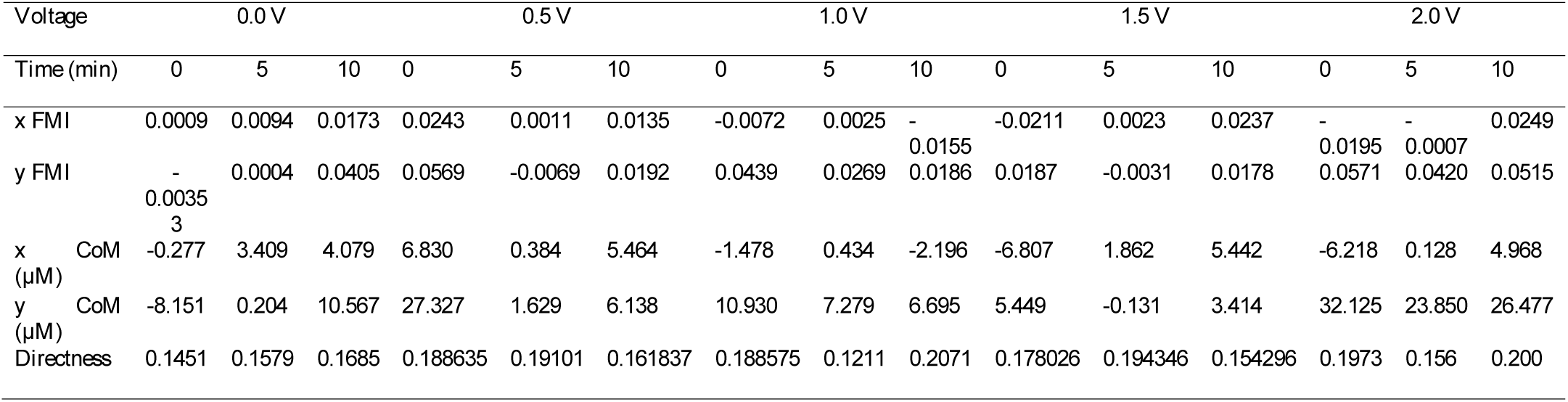
Directness, FMI and COM to ensure proper trials were conducted. The directness significantly increased from by the results of microchannel geometry due to the applied electric field. Values are mean based on a population of 30 cells for 60 second tracking.

Both successive and sudden voltage application methods were used to analyze the response of *P. aeruginosa* to electric fields. The sudden voltages applied were compared against successive voltages of the same magnitude at t = 0 min (Table 3). There were small changes in cell motility with different voltage application methods. Successive voltage applications showed a greater reduction in the accumulated distance and velocity compared to the sudden voltage applications. The distances travelled by the bacterial cells were 17.57 and 16.33 µm/s at the successive voltages of 1.0 and 2.0 respectively, while the distances were 18.58 and 18.85 µm/s for the sudden voltages of 1.0 and 2.0. When comparing the directness between the two application methods (Fig. 5D), we found that prolonged exposure to applied voltage further increases the trajectory of the cell. The accuracies with successive voltage applications were higher at 0.1886 (1.0 V application) and 0.1974 (2.0 V application), compared to 0.1581(1.0 V) and 0.1436 (2.0 V) with sudden voltage applications. Statistical significance was determined using a 1 way ANOVA with Dunnett’s post hoc test with p<0.05 (Table 4).

**TABLE 3.**
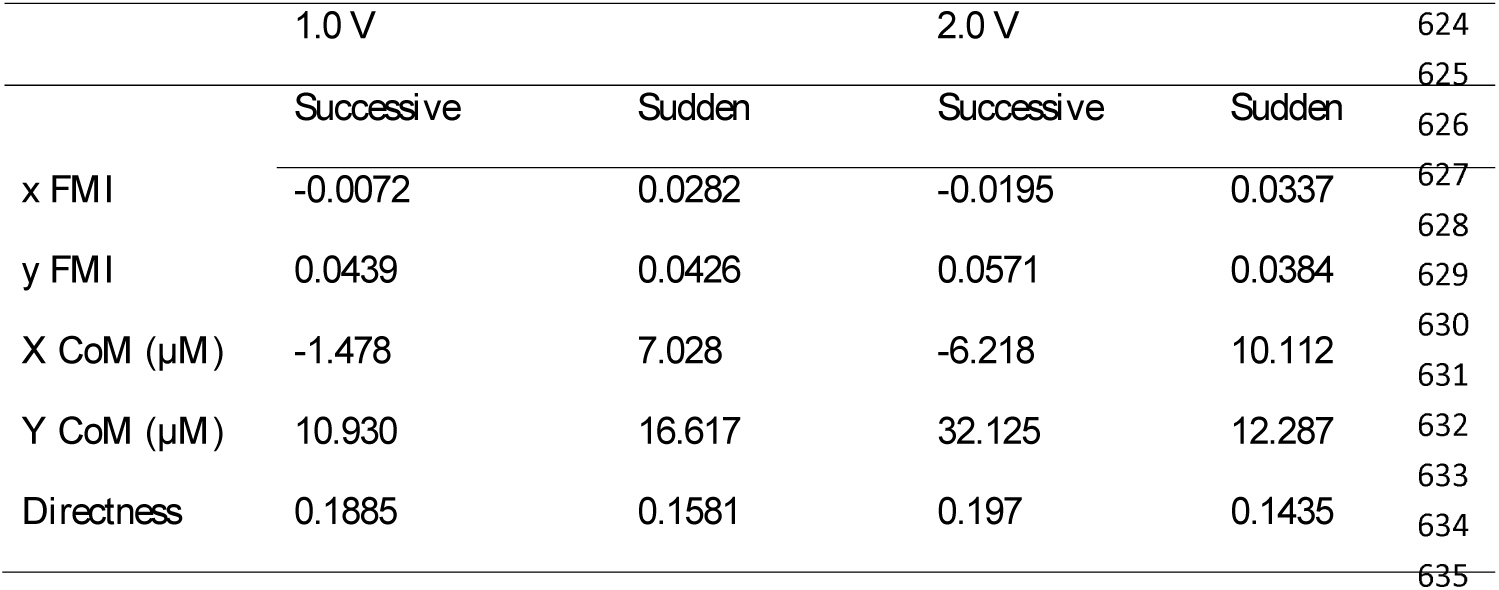
Comparisons of sudden and successive voltage application methods. Directness change is notably different between the two methods. Values are mean based on a population of 30 cells for 60 second tracking.

|  | 1.0 V |  | 2.0 V |  | 624 |
| --- | --- | --- | --- | --- | --- |
|  | Successive | Sudden | Successive | Sudden | 625 |
| x FMI | -0.0072 | 0.0282 | -0.0195 | 0.0337 | 626 |
| y FMI | 0.0439 | 0.0426 | 0.0571 | 0.0384 | 627 |
| X CoM (μM) | -1.478 | 7.028 | -6.218 | 10.112 | 628 |
| Y CoM (μM) | 10.930 | 16.617 | 32.125 | 12.287 | 629 |
| Directness | 0.1885 | 0.1581 | 0.197 | 0.1435 | 630 |
|  |  |  |  |  | 631 |
|  |  |  |  |  | 632 |
|  |  |  |  |  | 633 |
|  |  |  |  |  | 634 |
|  |  |  |  |  | 635 |

**TABLE 4.** Representative ANOVA results on the trend of motility factors observed for microchannel geometry effect, elecrotaxis and durotaxis gradient experiments. A value of p< 0.05 was considered statistically significant.

| Treatment | Parameter | Degrees of freedom | F | P |
| --- | --- | --- | --- | --- |
| **Microchannel geometry devices** |  |  |  |  |
|  |  | 2 | 97.54 | 0.00 |
| Circular | Accumulated distance (between groups) | 2 | 3.40 | 0.016 |
| Semi-circular |  | 2 | 163.75 | 0.00 |
| Square-tooth | Euclidean distance (between groups) |  |  |  |
|  | Cell velocity (between groups) |  |  |  |
| **Durotaxis** |  |  |  |  |
| Control (no coating) |  |  |  |  |
| 0.75 μg/ml collagen coated surface | Migrated distance across the channel | 2 | 480.93 | 0.00 |
| 150 μg/ml collagen coated surface |  |  |  |  |
| **Electrotaxis** |  |  |  |  |
| Successive voltage 0.0 V | Accumulated Distance (between groups) | 3 | 283.6 | 0.0265 |
| Successive voltage 1.0 V |  | 3 | 829.43 | 0.00 |
| Successive voltage 2.0 V | Euclidean Distance (between groups) | 2 | 8.859 | 0.003 |
|  | Cell velocity (between groups) |  |  |  |

## Discussion

Cellular migration is the result of the cumulative impact of a combination of multi-cue environmental inputs. Within a wound matrix, there is a high degree of spatial heterogeneity and topographical differences that are directly responsible for directed cell movement. The geometry and directional variation of the micro-deformations guide an individual bacterium in either a uniform manner for isotropic deformations or a random manner for anisotropic deformations (Cohen et al., 2014). Cell directness is impacted by the relevant surface features present at a wound site, as supported by our findings that chamber geometry impacts both directness and velocity. Similar observations were reported in a previous study by Doyle and colleagues (Doyle et al., 2009), where directness was attributed to the anisotropy of the pillar arrangement, and the corresponding centre of mass. *P. aeruginosa* motility characteristics are dependent on use of flagella and/or pili, expression of which is influenced by the external environment. The hydrodynamic streamlines of flow around the geometries present in the microfluidic chamber also influence cell motility, which can be described by rheotaxis, as bacteria adjust to the shear forces present in the medium (Marcos et al., 2012). Mechanical deformations specific to different geometries present along with the micro-flow environment along with the variation and presence of aqueous buffers influence the choice of motility and encourage bacteria to implement multiple motility mechanisms.

Durotactic cell migration studies shows that eukaryotic cells are attracted to an environment of increased stiffness (Lara and Schneider, 2013). Typical experiments have demonstrated that durotactic migration occurs with polymers, such as polyacrylamide, although little is known about the response to natural wound matrix proteins, such as collagen. We found that individual cell motility showed an attraction towards environments of increasing stiffness, as the concentration of type-1 collagen was increased under directed cell movement (20 μL/h flow rate). Our study is inherently different than many durotactic trials, where group cellular movement is observed, followed by the quantification of biofilm formation after a set growth period. An accepted mechanism for the observed *Pseudomonas aeruginosa* behaviour is through the regulation and adjustment of the composition and stiffness of the surrounding matrix (Kidoaki, 2010). Through this mechanism, combined with changes in cytoskeleton orientation, cell migration and growth are drawn towards matrices with higher stiffness (Kidoaki, 2010). Multiple proteins and adhesion reaction complexes within the cytoskeleton are responsible for directed cell movement (Kidoaki, 2010). For our initial study, we have investigated only collagen (as proof of principle), but we plan to expand future studies to examine the impact of other relevant matrix proteins (e.g. hyaluronic acid).

Because the impact of electrical current on bacterial motility has received limited attention, the rationale for the choice of specific times and applied voltage in this study was based on the need to cover a broad range to get a better idea of what variables to examine and how they influence motility. A range of 0 to 2 V produced electric field strength strong enough at low current that may exist in a wound matrix (Messerli and Graham, 2011). The strength and the time of exposure of the electric field were chosen based on the desired field strength inside the microenvironment surrounding the bacterial cells in the microfluidic chamber. The voltage drop caused by the resistance in the flow ports of the chamber; the platinum electrode and the buffer solution were considered when deciding the ranges of the applied electric field. Cell directness was enhanced by voltage compared to the microchannel geometry trials where no electrical field was applied.

Electrotaxis may open up promising applications to inhibit cell proliferation and bacterial growth in mammalian cell cultures. Directed cell movement occurs in response to an electrical potential gradient, but is specific to individual cell species (Cohen et al., 2014). *Pseudomonas aeruginosa* cells were attracted to the cathode in our experimental setup but this cannot be generalized across additional species or within polymicrobial biofilms (Lara and Schneider, 2013; Doyle et al., 2009). The electrode to which the Dictyostelium eukaryotic cells are attracted may depend on the proteins expressed during the time of electrical stimulation. Additionally, concentrations of Ca^2+^ and extracellular K^+^ ions can cause the cellular membrane to depolarize or hyperpolarize, depending on its relative orientation in respect to the anode and cathode (Gao et al., 2011). The membrane voltage facing the anode is hyperpolarized and attracts Ca^2+^ ions, resulting in a contraction (Mycielska and Djamgoz, 2004). The resulting contraction causes movement towards the cathode, depolarizing the opposite membrane and opening the voltage-gated Ca^2+^ channel, eventually resulting in additional ion influx and propulsion (Gao et al., 2011) in the opposite direction. Overall cell motility is further dependent on membrane voltages, exterior K^+^ concentrations, and Ca^2+^ balance (Gao et al., 2011). While typical wound matrix electrical fields can range from 42 to 100 mV mm^−1^, the studied voltages were applied in a range up to 2.0 V for sudden or successive application methods to understand the possibilities of using similar voltages to limit cell velocity and/or control directionality. The velocity of single *P. aeruginosa* cells decreased with exposure to increasing voltages for prolonged periods of time, up to 10 min in length. This observed effect is distinctly different from the response seen in mammalian cells.

Studies have shown that the velocity in eukaryotic cells increases with the application of voltage, which is in direct contrast to the results obtained with *P. aeruginosa* in our present study (Hart et al., 2013). Endogenous electrical signals represent an important signalling mechanism that guides cell migration during wound healing, ultimately regulating the direction of cell movement. Therefore, the application of voltage may have specific influences on bacterial versus eukaryotic cells and could represent a specific therapeutic application for the treatment of bacterial biofilms occurring on or within human tissues. However, additional studies are necessary to determine the overall impact of voltage application on biofilm formation, infectivity, wound pathology, and healing outcomes. In the course of our work, in keeping with other findings, we have demonstrated that microfluidic devices may be an ideal tool for studying the electrotaxis characteristics of single bacterial cells, which may aid in the identification of new therapeutic avenues for the treatment of recalcitrant biofilm infections, specifically those that slow or prevent wound healing (Li and Lin, 2011).

## Author Contributions

SN conceived the study. SN and LP designed the experiments. LP conducted the experiments with assistance from SN. LP and SN analyzed the data. LP wrote the manuscript with input from SN. All authors read and approve the manuscript.

## Acknowledgments

The authors sincerely thank the Natural Sciences and Engineering Research Council of Canada, and the Ontario Ministry of Research and Innovation for funding this study.

